# Assessing the Live Decellularized Renal Microvasculature Post-Transplantation Using Intravital Microscopy

**DOI:** 10.1101/2021.10.07.463473

**Authors:** Peter R. Corridon, Anousha A. Khan

**Author notes:** **Corresponding Author:** Peter R. Corridon, PhD, Assistant Professor, Department of Immunology and Physiology, College of Medicine and Health Sciences, Director, Pre-Medicine Bridge Program, College of Arts and Sciences, Khalifa University of Science and Technology, PO Box 127788, Abu Dhabi, UAE, Office.

## Abstract

Transplantation is the ideal solution for end-stage renal failure, but the growing mismatch between organ supply and demand accentuates the need for alternative solutions like the bioartificial kidney. Several approaches to developing this technology have been demonstrated, and whole organ decellularization appears to be a promising methodology. One major challenge to this strategy is maintaining vascular integrity and functionality post-transplantation. Most models to examine the microvasculature have primarily utilized in vitro or in vivo techniques that are incapable of providing adequate spatial and temporal resolution. Here, we show that decellularized scaffolds orthotopically transplanted into rats initially retain microvascular structure in vivo using intravital two-photon microscopy, as previously identified in vitro. Large molecular weight dextran molecules also provide real-time evidence of the onset of ischemia and increases in microvascular permeability, support substantial translocation of dextran macromolecules from glomerular and peritubular capillary tracks as early as 12 hours after transplantation. Macromolecular extravasation continued across a week, at which time the decellularized microarchitecture was significantly compromised. These results indicate that a in vivo method capable of tracking microvascular integrity represents a powerful interdisciplinary approach for studying scaffold viability and identifying ways to promote scaffold longevity and angiogenesis in bioartificial organs.

## Introduction

Although limited by the scarcity of organs, transplantation is the ideal solution for end-stage renal disease (ESRD), mainly due to the limited nature of dialysis^1, 2^. Characterized by a debilitating and irreversible loss of renal function, the exponential increase in ESRD has accentuated the demand for alternative solutions like the bioartificial kidney^3, 4^. One of the most exciting approaches for developing a bioartificial kidney is whole organ decellularization^5^. Decellularization is a promising bioengineering strategy because it focuses on extracting the extracellular matrix (ECM) from the native kidney, ideally with intact structural and functional cues. The ECM can then be used as a natural scaffold for recellularization. This interdisciplinary approach applies principles of life sciences and engineering toward the development of functional replacement organs.

The optimal conditions to achieve a decellularized whole organ need to be identified. Nevertheless, these conditions are dependent on the size, density, and complexity of the tissues. Organs can be decellularized by different methods, including but not limited to physical or chemical disruption, as well as enzymatic treatment via perfusion- or immersion-based techniques. Sodium dodecyl sulfate (SDS) is a critical component of the majority of decellularization protocols. This widely used ionic surfactant is capable of rapidly and efficiently removing cells and genetic material form native structures^5, 6^. Also, perfusion-based techniques are preferred, particularly for highly vascular organs like the kidney, as they utilize the innate organ vasculature to deliver the decellularization agent providing faster and more homogenous acellular templates^7^.

Several microscopic and macroscopic techniques have been employed to examine whole kidney scaffold structure and function in various settings^8-15^. Such investigations have provided evidence that the decellularization process successfully removes remnant native cellular/tissue components and decellularizing agents, while maintaining the vascular and ECM architectures in various large and small animal models. Preserving the vascular architecture is essential as it provides the ideal template for cellular repopulation^16^. Moreover, the effective removal of scaffold remnants is crucial for inhibiting detrimental biological and immunological effects during recellularization and after transplantation.

Beyond these achievements, regrettably, there are still events that considerably compromise vascular integrity and functionality. These events are observed essentially in a post-transplantation setting, regardless of the decellularized or recellularized nature of the scaffold. The combination of micro-and macro-vascular complications observed with current renal transplantation regimens is far more likely to occur in scaffolds with altered structural integrities. Most models used to examine the decellularized renal vasculature have primarily utilized in vitro or in vivo imaging techniques to investigate events at the microvascular level, namely alterations in endothelial and epithelial layer porosity. However, these methods cannot provide adequate spatial and temporal resolutions, and such limitations outline the need for alternative ways to explore these events.

Fortunately, intravital multiphoton microscopy offers a unique opportunity, as it offers subcellular resolution to characterize live morphological and functional features within the kidney^17-20^. As a result, this study aimed to provide a real-time in vivo assessment of the microarchitecture of the decellularized kidney post-transplantation. To achieve this, techniques from previously optimized protocols that preserve the inherent vascular network and limit cell toxicity were applied^12, 21^. Whole kidneys were extracted from rats, decellularized by perfusion of 0.5% SDS for roughly 8 hours. These scaffolds were then sterilized and orthotopically transplanted into their respective donors. After that, a combination of nuclear and vascular dyes (Hoechst 33342 and 150-kDa FITC dextrans, respectively) provided real-time evidence of the effective removal of cell/nuclear remnants from the acellular scaffolds. Drastic changes in glomerular/peritubular capillary permeability, vascular flow, and blood extravasation accompanied ischemia/thrombosis during the week that followed transplantation. This advanced imaging technique provides submicron spatial resolution, and millisecond temporal resolution^18^, which are ideal for visualizing live cellular/subcellular events in real-time. Even though specialized, intravital microscopy has shown great utility in provide real-time assessment of renal structure and function in vivo and can thus open new horizons for the detailed evaluation of bioartificial organs.

## Results and Discussion

### Biochemical Assays Used to Evaluated the Decellularization Process

A protocol based on the primary use of a single ionic surfactant was employed to decellularize native rat kidneys. As reported in the literature^12, 22^, the 0.4 ml/min infusion of only SDS via a cannulated renal artery extensively removed cellular materials from the native kidneys. Substantial visible changes to the original organ (Figure 1A) occurred as early as 4 hours (Figure 1B), as it ultimately transitioned to a translucent structure (Figure 1C). The resulting scaffolds were then perfused with PBS to remove the cellular and detergent remnants (Figure 1D) and subjected to biochemical assays to evaluate residual DNA (Figure 1E) and SDS (Figure 1F). The assays revealed that this method removed approximately 98% of the innate DNA content from original kidneys and 99% of the remnant SDS from the scaffold. The significant removal of DNA (Kruskal-Wallis, p < 0.001) and SDS (Kruskal-Wallis, p < 0.001) confirmed the effectiveness of this protocol.

**Figure 1.**
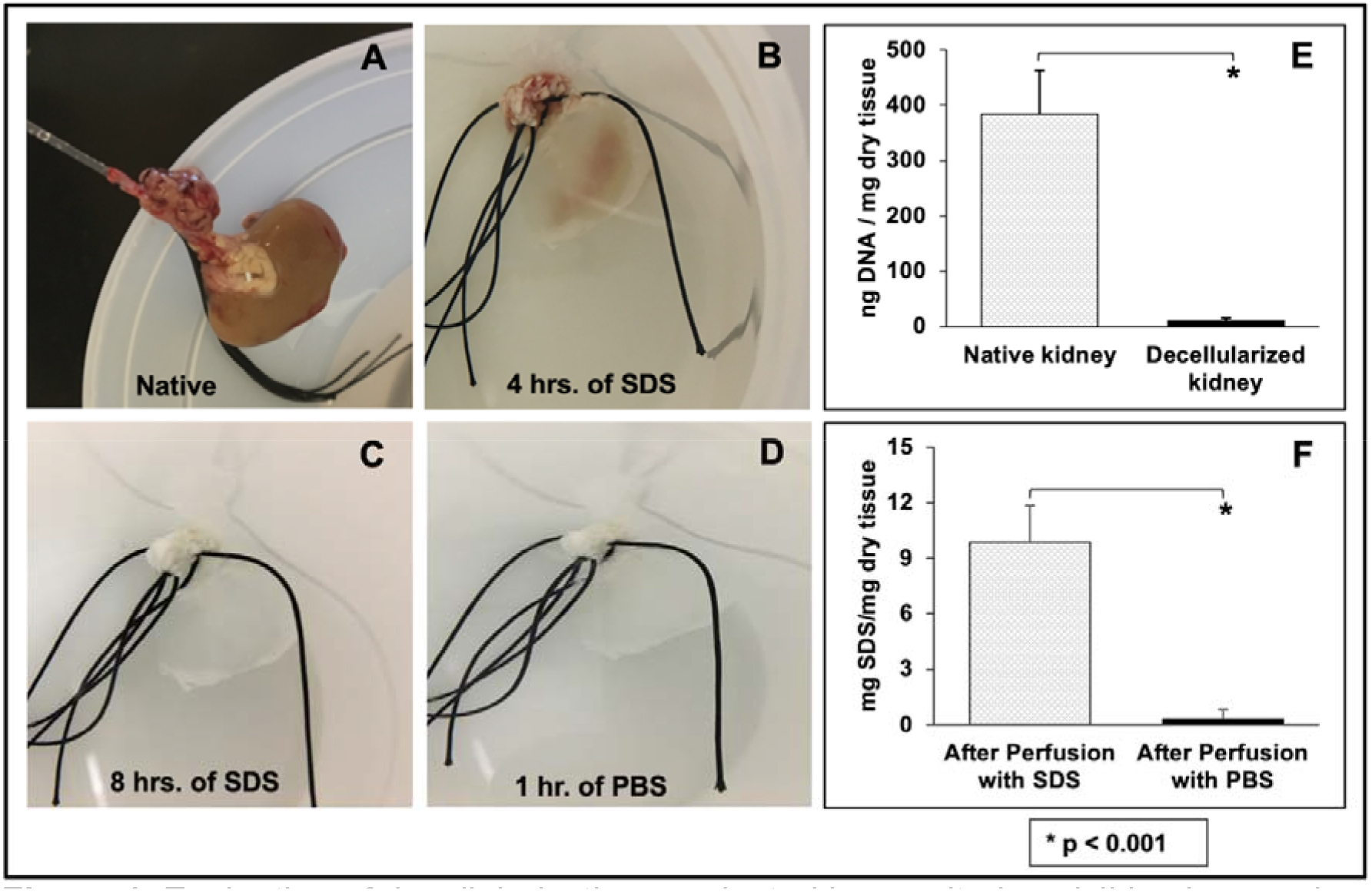
Evaluation of decellularization conducted by monitoring visible changes in the organ and biochemical assays used to measure residual DNA and SDS concentrations. A) An image of harvested whole rat kidney with its renal artery cannulated for perfusion-based decellularization. B) The kidney after 4 hours of perfusion with SDS illustrates its transition from solid to translucent. C) The fully decellularized kidney was produced after 8 hours of SDS perfusion. D) The kidney after its subsequent perfusion with PBS. E) A plot presenting the low level of remnant DNA in the scaffold after decellularization. F) A graph highlighting the effective removal of SDS from the scaffold and its low residual concentration. Non-parametric evaluations conducted using Kruskal-Wallis detected significant declines in both DNA and SDS concentrations after decellularization (* p < 0.001).

Evaluating the concentrations of these residual components is a crucial aspect of the decellularization process. It is necessary to efficiently remove cellular remnants from scaffolds to reduce the potential for unwanted immunological responses or tissue rejection after transplantation^23^. It is also essential that residual concentrations of SDS be low enough so that the detergent does not continue to denature the scaffold. This unwanted effect can alter the scaffold’s overall permeability, and compromise its integrity, which can altogether adversely affect recellularization and transplantation^24^.

### Whole Rat Kidney Decellularization Evaluated by Intravital Two-Photon Fluorescence Microscopy

As proof of principle, intravital two-photon microscopy was used to examine the effectiveness of the decellularization process further. Live blue pseudo-color channel images obtained revealed considerable variations in fluorescence from native (non-transplanted) kidneys (Figure 2A) and decellularized scaffolds that were transplanted into live recipients after introducing Hoechst 33342 (Figure 2B). This membrane permeant dye diffuses readily into tissues^25-30^, where it then labels DNA in live and fixed cells by binding to adenine-thymine-rich regions of DNA in the minor groove to produce large enhancements in fluorescence^31^. Generally, such enhancements are known to support the clear visualization of nuclei in the endothelia, epithelia, glomeruli, interstitial cells, and circulating leukocytes ^28^.

**Figure 2.**
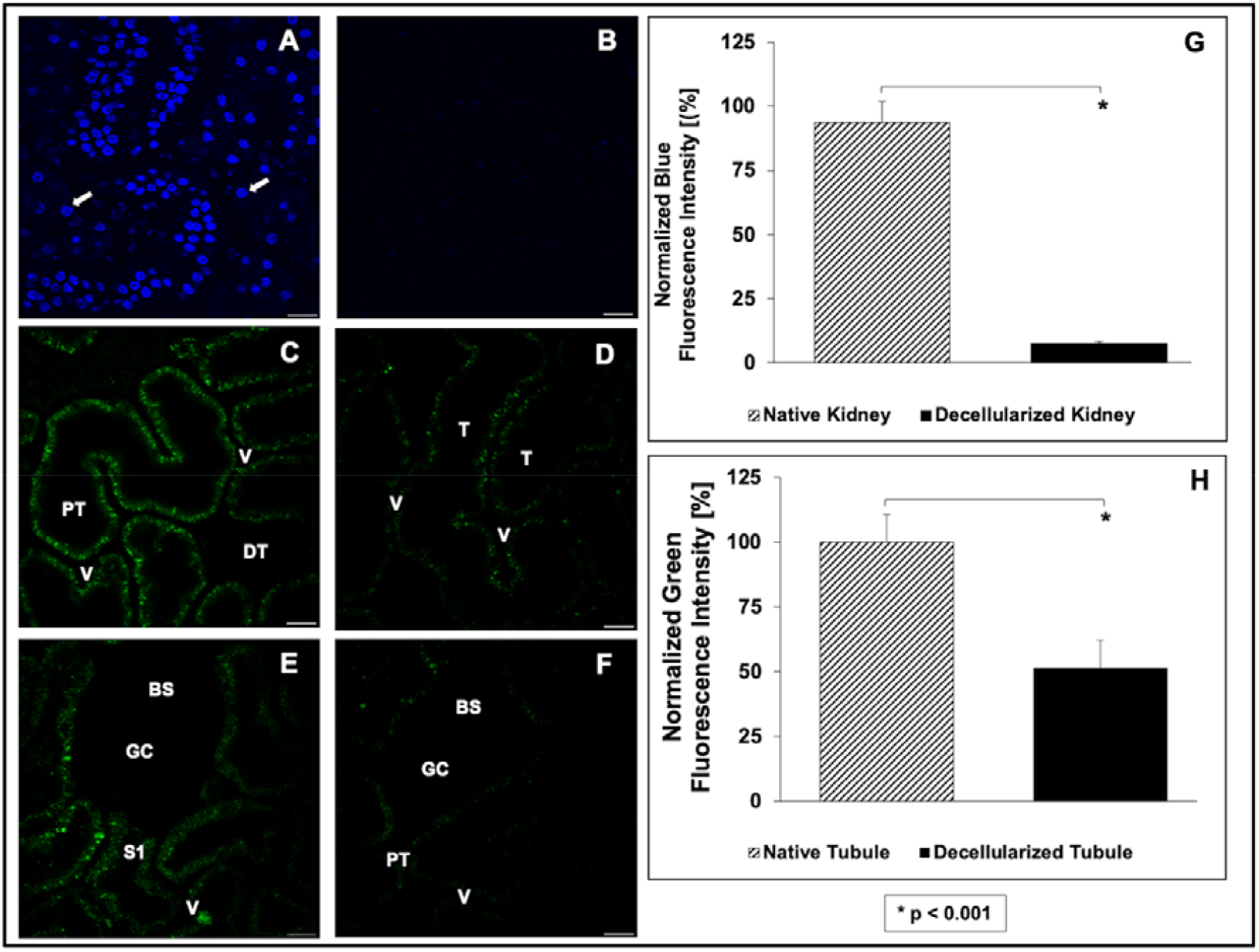
Whole rat kidney decellularization evaluated by intravital two-photon microscopy using Hoechst 33342-based fluorescence and innate autofluorescence. A comparison of blue pseudo-color channel images illustrating Hoechst 33342 dye fluorescence intensity signals. A) Images obtained from native (non-transplanted) showing the vibrant presence of the nuclear stain. B) Image from a transplanted decellularized kidney showing the absence of Hoeschest labeling. A comparison of green autofluorescence signals gathered from native (non-transplanted) and transplanted decellularized kidneys. C) Image obtained from a native kidney that presents only distal (DT) and proximal (PT) tubular compartments and peritubular vascular tracks (V). D) Intravital micrograph identifying decellularized tubular (T) and vascular compartments. It should be noted that the decellularization process made it difficult to differentiate between tubular segments. E) Image obtained from a native kidney that captured the Bowman’s space (BS) and glomerular capillaries (GC), the S1 segment of the proximal tubule compartments (S1), and peritubular vasculature. F) Image of a decellularized kidney highlighted the decellularized glomerular and tubular segments. Scale bars represent 20 µm. Non-parametric evaluations conducted using the Kruskal-Wallis test detected significant reductions in both the normalized blue (* p < 0.001) and green (** p < 0.001) pseudo-color fluorescence observed after decellularization.

Fluorescence intensity measurements obtained from these images showed an approximate 92% drop in the relative level of fluorescence in the decellularized scaffold and the absence of nuclear staining compared to the native kidney. This significant reduction in the relative blue pseudo-color fluorescence (Kruskal-Wallis, p < 0.001) correlates with the decrease in DNA content obtained from the biochemical assay (Figure 1E). These in vivo results also complement conventional in vitro histological and fluorescent microscopic methods that have been routinely applied to investigate decellularization^32^.

Similarly, images collected from the green pseudo-color channel displayed variations in the intrinsic level of autofluorescence present in native kidneys (Figure 2C and Figure 2E) compared to that of the decellularized organ (Figure 2D and Figure 2F). Normally, the kidney has a high level of green autofluorescence^25, 26^, characterized by the natural emission of light by biological structures such as mitochondria and lysosomes^20^, and unique metabolites like aromatic amino acids, nicotinamide adenine dinucleotide (NADH and its phosphate analog NADPH), and flavins^33, 34^. The relative distribution and percentages of these structures vary and help distinguish renal compartments, namely the proximal and distal tubules, glomerulus, peritubular capillaries, and interstitium, without the addition of fluorescent markers^9, 28^.

Among these structures, proximal tubules are easily identified as having the greatest autofluorescent signal and thickness, whereas distal tubules appear thinner and dimmer (Figure 2C). Likewise, the characteristic shape of the renal corpuscle is highlighted by the outline of the Bowman’s capsule and faint or invisible capillary tuft^35^, along with the nearby peritubular capillary and interstitial space (Figure 2E). However, after decellularization, significant (Kruskal-Wallis, p < 0.001) reductions in the green pseudo-color signal intensity were observed. Specifically, there was an approximate 50% drop in the relative level of fluorescence in the scaffolds (Figure 2H), which made it difficult to identify specific tubular compartments (Figure 2D), except proximal tubule segments that emanated from the Bowmans capsule (Figure 2F). Overall, similar drastic changes in tissue fluorescence were also previously observed using immunohistochemistry approaches in vitro and confirm decellularization^12, 24, 36^.

### Real-time In Vivo Examination Minor Alterations to Blood Flow and Dextran Leakage within Scaffolds Directly After Transplantation

Once decellularization was established, intravital microscopy was then utilized to examine the structural and functional integrity of the transplanted scaffolds in vivo. Scaffold integrity was first investigated immediately post-transplantation, whereby time-series data was used to monitor the live infusion of the 150-kDa FITC dextran. This dextran is a branched polysaccharide. The fluorescein moiety is attached by a stable thiocarbamoyl linkage, which does not lead to any depolymerization of the polysaccharide and helps the dextran retain its minimal charge. This feature is an essential requirement for permeability studies.

In vivo data showed that the fluorescence levels in the tubular epithelial and luminal compartments remained relatively constant and comparable in both native (Figures 3A-3C and 3G) and decellularized (Figures 3D-3F and 3H) kidneys. In comparison, fluorescence intensity levels within the vascular lumens of native and decellularized kidneys were substantially higher than the previously mentioned other regions directly after transplantation. The Kruskal-Wallis test revealed that these differences within native kidneys were significant (p < 0.001), and the post hoc Dunn’s test only detected significant pairwise differences between the fluorescence intensity in the vascular lumen and the tubular epithelium (p < 0.001), and vascular lumen and the tubular lumen (p < 0.001). Likewise, for decellularized kidneys, this non-parametric test also revealed a significant difference among the intensities in the tubular epithelia, tubular lumen, and vascular lumen (p < 0.001). Thereafter, the post hoc test only detected significant pairwise differences between the fluorescence intensity in the vascular lumen and the tubular epithelium (p < 0.01), vascular lumen and the tubular lumen (p <0.001), and tubular epithelium and the tubular lumen (p = 0.02).

**Figure 3.**
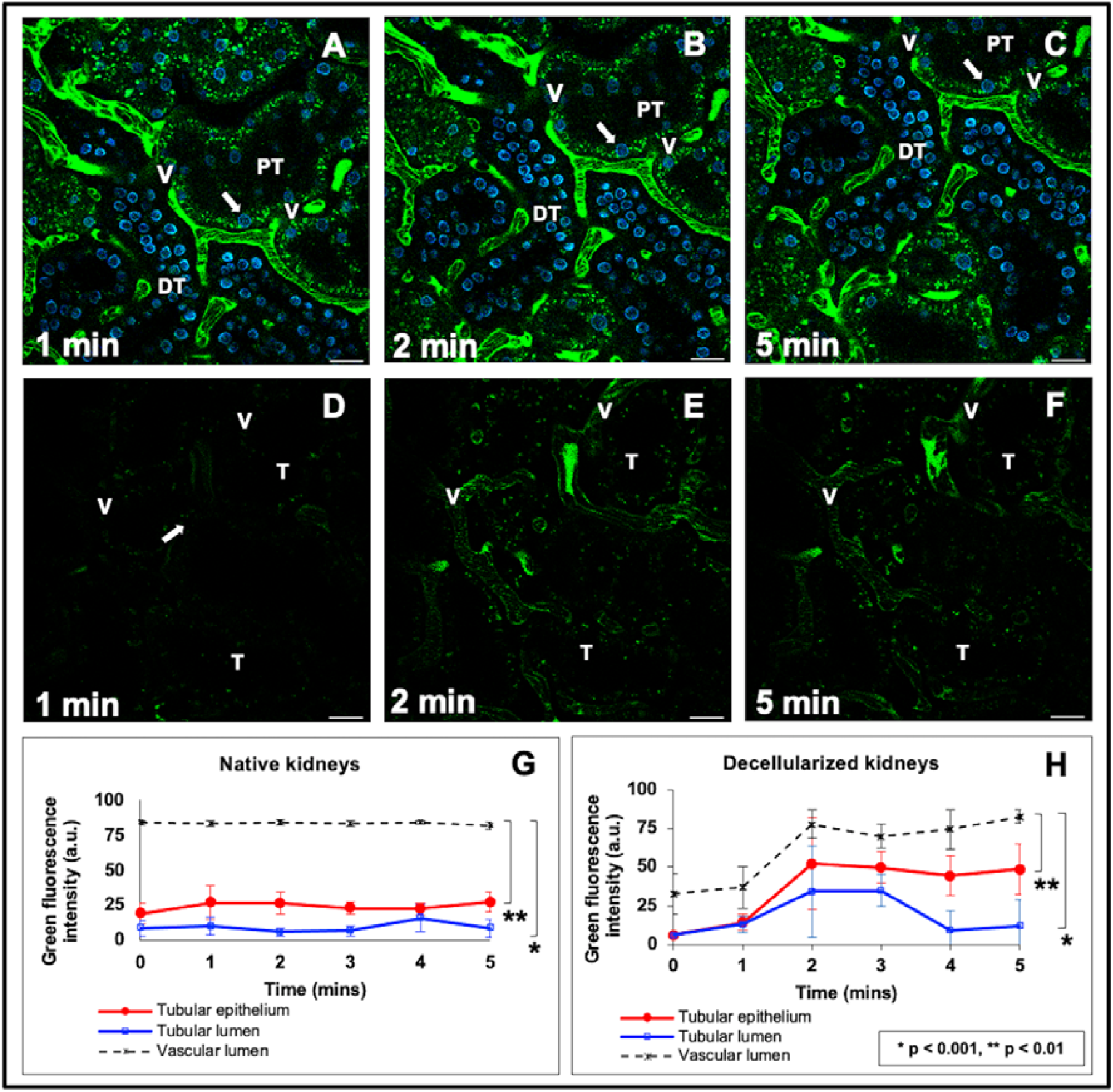
An in vivo assessment of the integrity of the decellularized rat kidney conducted immediately after transplantation using 150-kDa dextran molecules. A) through C) Time-series images taken at 1 min, 2 min, and 5 min marks from a live native kidney display the proper confinement of the large molecular weight dextrans within the peritubular vasculature (V) and characteristic autofluorescence levels within proximal (PT) and distal (DT) tubules, and nuclear staining with Hoechst 33422 (arrows). D) through F) Images obtained at 1 min, 2 min, and 5 min marks from a transplanted acellular scaffold also display the confinement of the large molecular weight dextrans within the peritubular vasculature (V) and substantially reduced autofluorescence levels within proximal (PT) and distal (DT) tubules, and absence of nuclear staining. Scale bars represent 20 µm. G) A graphical representation of the variations in fluorescence levels (obtained from the green pseudo-color channel) associated with the time-series data for native kidneys outlined the relatively constant and elevated level of fluorescence within the vasculature, compared to the much lower signals recorded in the tubular lumen and epithelium. H) A similar graphical representation of the variations in fluorescence levels (obtained from the green pseudo-color channel) associated with the time-series data for decellularized kidneys outlined the rising level and FITC fluorescent within the vasculature, compared to the much lower signals recorded in the tubular lumen and epithelium. The data also suggests some minor degree of dextran leakage from the microvasculature across this period. It should be noted that there were minor shifts in the field during image acquisition that resulted from vibrations caused by respiration. Time-lapse video showing portions of these event are presented in Supplemental Video 1 (native kidney) and Supplemental Video 2 (decellularized kidney).

During the immediate timeframe after transplantation, the dextran molecules were exclusively present in vascular lumens. However, there was a delayed, lower, and inhomogeneous distribution of the dextran observed in the decellularized vasculature (Figures 3D-3F and 3H) compared to the more homogenous distribution of the FITC dye in normal kidneys imaged under the same conditions (Figures 3A-3C and 3G), as more time was possibly needed to fill the acellular nephron with blood directly after transplantation. Our statistical analyses also indicated that the variations in quantity of blood within native and decellularized kidneys were significantly different within this period (p = 0.034). The quantity of blood was indirectly estimated by the level of dextran fluorescence intensity within the microvascular lumen.

Overall, the data provided real-time evidence that these large molecular weight molecules were primarily confined to the decellularized vascular lumen directly after transplantation. However, the data also suggested that the dye began to leak through the modified renal filtration and peritubular barriers (Figure 3D through Figure 3F, Supplemental Video 2) in comparison to native, non-transplanted kidneys (Figure 3A through Figure 3C). This view is based on the relative significant variations in fluorescence levels observed in the tubular epithelial of the native and decellularized kidneys during this period (p < 0.001), as well as the non-statistically significant difference between signals observed in the native and decellularized lumens (p = 0.057). Such results emphasize the power of intravital microscopy to uncover subtle changes in vascular permeability, even with the use of a small sample size. Moreover, the explanted decellularized kidney vasculature was able to withstand some degree of in vivo blood flow/pressure levels initially, as no substantial signs of rupturing were detected at the macroscopic or microscopic level.

### Severe Dextran Extravasation and Alterations to Capillary Blood Flow Observed Across One Week of Transplantation

Vascular permeability was further investigated throughout the week that followed transplantation. To support this investigation, the 150-kDa FITC dextran was infused into the tail vein directly before blood was introduced into transplanted scaffolds. This process provided a means to evaluate the distribution of the blood constituents and insight into the in vivo environment’s effect on the grafts. This infusion regimen was favored over infusing the dye at later times, as it was necessary to ensure that the fluorescent marker circulated through the nephron because, at future time points, clotting could prevent the dye from entering the microvasculature.

With this approach, drastic changes in vascular structure and function from the initial state (Figure 4A) were detected at the 12-hour (Figure 4B), 24-hour (Figure 4C), and 1-week (Figure 4D) time points. The varying degrees of dextran extravasation from the microvasculature observed across this period signify the impairment of typical filtrative capacities and increased permeability of the decellularized microvasculature under in vivo conditions. Normally, the kidney can autoregulate blood flow to safeguard against considerable fluctuations in blood pressure transmitted to peritubular and glomerular capillaries^37^. However, the acellular organ is incapable of replicating this vital function^9^, and as a result, scaffolds would have been exposed to pressures capable of damaging these delicate structures.

**Figure 4.**
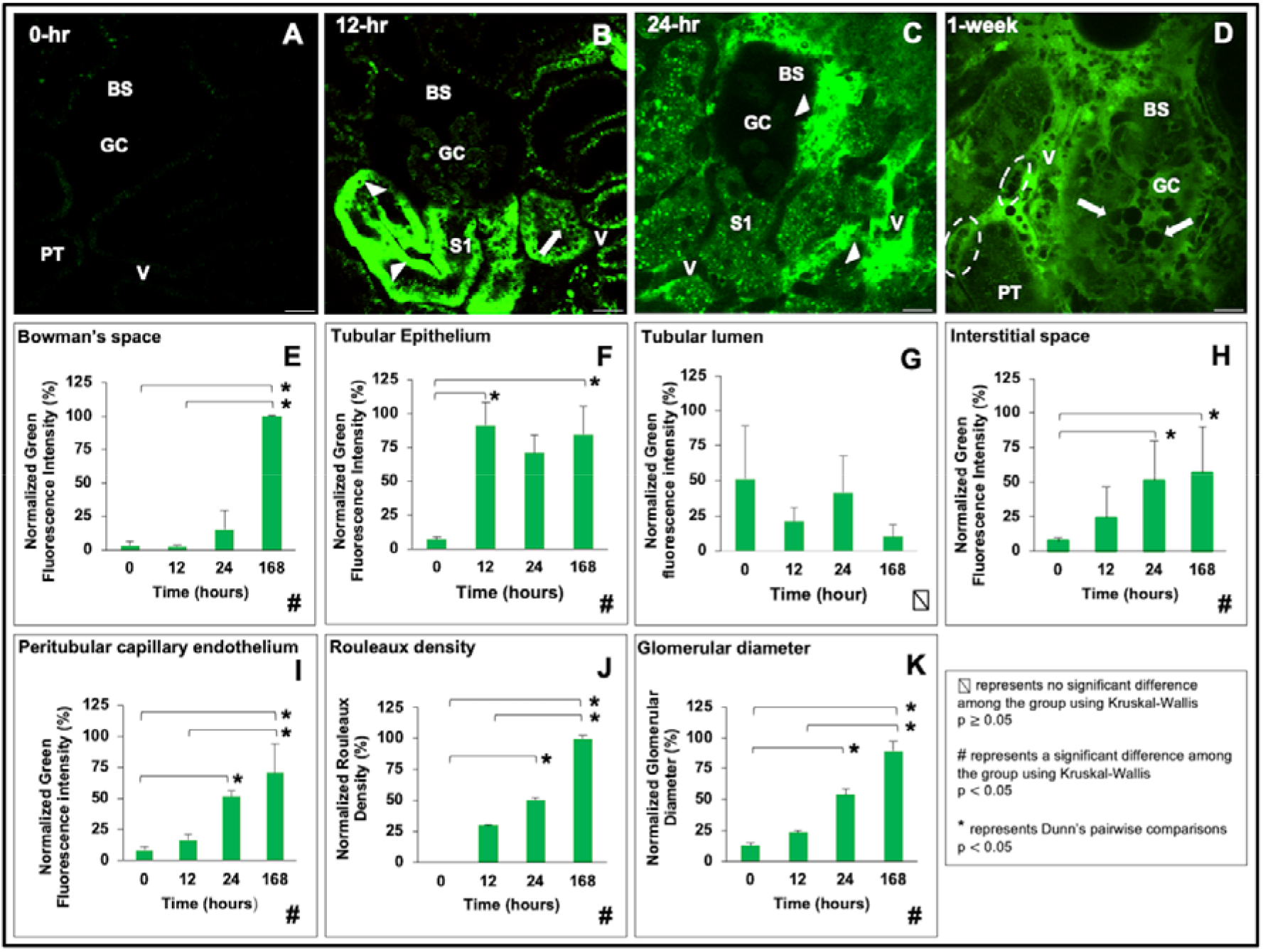
Impact of the in vivo environment on scaffold integrity during a week after transplantation. A) Image taken from a decellularized kidney that displays the decellularized autofluorescence before the introduction of FITC (this is the same imaging field that is shown in Figure 3E). B) Image taken from a decellularized kidney 12 hours after transplantation illustrates substantial and inhomogeneous levels of dye translocation between luminal, epithelial, and interstitial compartments. C) Image taken from a decellularized kidney 24 hours after transplantation presents the accumulation of the dextran and blebs within the Bowman’s capsule, interstitium, and tubules. D) Image taken from a transplanted decellularized kidney 1 week (168 hours) after transplantation provides evidence of significant rouleaux (dashed oval within the vasculature) and bleb/vesicle (arrows within the Bowman’s space and tubular lumen) formation that accompanied dye extrusion from breached glomerular capillaries to completely occlude this enlarged glomerulus. Scale bars represent 20 µm. Graphs examining the degree of FITC dye translocated within the E) the Bowman’s space, F) tubular epithelium, G) tubular lumen, H) interstitial space, I) peritubular capillary endothelium, as well as J) rouleaux density, and K) glomerular diameter, during the 168-hour measurement period are accompanied by H) changes in rouleaux density.

Mechanistically, such damage would explain the progressive leakage of the FITC dye from the glomerular capillaries into the Bowman’s space (Figure 5). Decellularization would have advertently altered microarchitectural permeability, thereby disrupting the glomerulus’ ability to act as a natural sieve that limits the passage of only water and small solutes into the filtrate. Within the native glomerulus, the fenestrated capillary endothelial barrier restricts the passage of molecules smaller larger than 70 nM; the basement membrane, with its negative charge, restricts the passage of particles less than 1 kDa and favors the filtration of cations^38^. Foot processes on podocytes provide an additional size selectivity, by limiting the passage of molecules larger than 14 nM^38^. By removing this intrinsic selectively permeable barrier, through decellularization, macromolecular transport was no longer limited by size, shape, charge, and deformability^39^.

**Figure 5.**
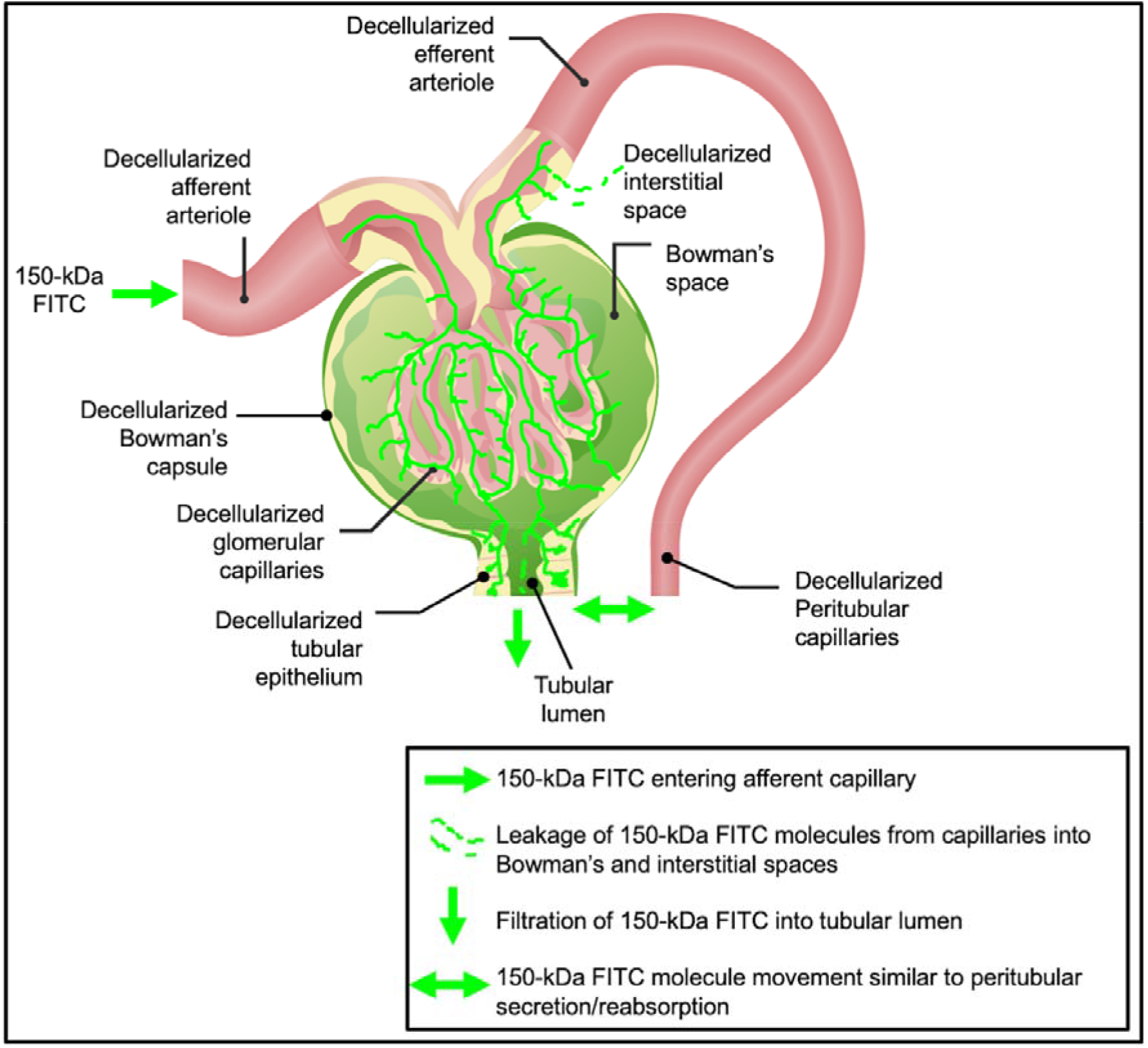
A schematic of possible mechanistic view of dextran extravasation from the decellularized vasculature and translocation to the decellularized peritubular epithelium and surrounding interstitial space. This model presents the considerable molecular weight, 150-kDa FITC, dye’s entry into the decellularized nephron vasculature via the afferent arteriole. The vascular marker then progresses through the decellularized glomerulus, where it can be filtered into the Bowman’s space and enter the tubular lumen. This unregulated process can potentially induce significant dynamic and static pressures to facilitate bilateral dye translocation between tubular epithelium, interstitium, and peritubular endothelium. The original image was adapted with permission^40^.

With the loss of the characteristics mentioned above, the acellular glomerulus would likely have facilitated the filtration of the 150-kDa molecules into the Bowman’s space and tubular lumen. The accumulation of blood within these regions would have created extravasation opportunities into the acellular tubular epithelium and interstitium. Thus, such translation would likely have mimicked unregulated secretion and reabsorption patterns throughout the fractured nephron. Specifically, after filtration, this dye would have subsequently entered the lumens of both the decellularized peritubular capillaries and tubular epithelium.

Furthermore, the simultaneous accumulation of fluid, macromolecules, and blood cells within these compartments could have established uncharacteristic hydrostatic, hydrodynamic, and osmotic pressure gradients and, in turn, supported the translocation into epithelial/interstitial compartments shown in the intravital micrographs (Figure 4A through Figure 4D). This effect could have undoubtedly reduced the concentration of the dye within the patent regions of the vasculature as evidenced within glomerular capillaries at the 12-hour and 1-week marks (Supplemental Video 3 and Supplemental Video 4, respectively). Correspondingly, from a quantitative perspective, measurements from these micrographs highlight one and two orders of magnitude increases in the presence of the dye within the Bowman’s space (Figure 4E), tubular epithelium (Figure 4F), tubular lumen (Figure 4G), interstitial space (Figure 4H), peritubular capillary endothelium (Figure 4I), rouleaux density (Figure 4J), and glomerular diameter (Figure 4K).

Likewise, alterations to capillary blood flow outlined by the increased presence of rouleaux were investigated post-transplantation. Rouleaux density measurements indicated one and two orders of magnitude increases in red blood cell aggregation (Figure 4J). Erythrocytic aggregation is a hallmark of ischemia^41^. This outcome was anticipated, as collagen within transplanted scaffolds would have been exposed to flowing blood. Such an interaction would have rapidly facilitated platelet activation and aggregation^42^. Within the ECM, collagen is the only protein that supports both platelet adhesion and complete activation^42^. Moreover, collagens in this matrix are normally separated from blood by the endothelial layer. However, the decellularized scaffold provides direct contact to flowing blood and effectively initiates hemostasis. It is expected that these processes were unregulated in the transplanted scaffolds and generated widespread coagulation and thrombosis, which would have ultimately facilitated the reductions in blood flow and entrapment of the dye within acellular nephrons observed at the 1-week mark.

These mechanisms are also known to support bleb and microvesicle formation visualized in vivo within decellularized glomerular segments. Bleb/microvesicle accumulation within the microvasculature can lead to longstanding occlusions, particularly in decellularized vessels that cannot compensate for blood pressure fluctuations. Furthermore, unresolved obstructions to blood flow generate detrimental stagnation pressures that can induce antegrade blood flow and overloaded renal compartments like the glomerulus. This process can lead to decellularized glomerular hypertrophy, an increase of diameter compared with that of the original, (Figure 4D and Supplemental Video 4). This form of hypertrophy could have arisen from considerable changes in efferent and afferent flows/pressures exerted on acellular glomeruli^43^. Excessive filtration and proteinuria could have also contributed to the condition^44^. Cellular debris and vascular cast formation in the Bowman’s space that could have arisen from ischemia-derived blood cell apoptosis or necrosis observed in chronic diseases are also likely contributing factors ^45^. Signs of these effects were observed by tracking the glomerular diameters, which markedly increased with time (Figure 4K).

Further analyses revealed statistically significant differences in the quantities of the vascular marker that were extruded from the capillaries across the 7-day measurement period using the non-parametric one-way ANOVA: Bowman’s space (Figure 4E, Kruskal-Wallis, p = 0.005); tubular epithelium (Figure 4F, Kruskal-Wallis, p = 0.014); interstitial space (Figure 4H, Kruskal-Wallis, p = 0.021); peritubular capillary endothelium (Figure 4H, Kruskal-Wallis, p = 0.004); rouleaux density (Figure 4J, Kruskal-Wallis, p = 0.005); and glomerular diameter (Figure 4K, Kruskal-Wallis, p = 0.004). Analogously, pairwise significant differences were detected at the following time point combinations shown in Table 1.

**Table 1.** Pairwise differences obtained using the post hoc Dunn’s test highlight the significant levels (p < 0.05) of FITC dye extravasation into various renal compartments across the week post-transplantation.

| Time point combination | Bowman's space | Tubular epithelium | Tubular lumen | Interstitial space | Peritubular capillary endothelium | Rouleaux density | Glomerular diameter |
| --- | --- | --- | --- | --- | --- | --- | --- |
| 0 – 12 | $p \geq 0.05$ | **$p = 0.002$** | $p \geq 0.05$ | $p \geq 0.05$ | $p \geq 0.05$ | $p \geq 0.05$ | $p \geq 0.05$ |
| 0 – 24 | $p \geq 0.05$ | $p \geq 0.05$ | $p \geq 0.05$ | **$p = 0.011$** | **$p \geq 0.009$** | **$p = 0.030$** | **$p = 0.017$** |
| 0 – 168 | **$p = 0.003$** | **$p = 0.014$** | $p \geq 0.05$ | **$p = 0.006$** | **$p = 0.001$** | **$p < 0.001$** | **$p < 0.001$** |
| 12 – 24 | $p \geq 0.05$ | $p \geq 0.05$ | $p \geq 0.05$ | $p \geq 0.05$ | $p \geq 0.05$ | $p \geq 0.05$ | $p \geq 0.05$ |
| 12 – 168 | **$p = 0.003$** | $p \geq 0.05$ | $p \geq 0.05$ | $p \geq 0.05$ | **$p = 0.030$** | **$p = 0.043$** | **$p = 0.017$** |
| 24 – 168 | $p \geq 0.05$ | $p \geq 0.05$ | $p \geq 0.05$ | $p \geq 0.05$ | $p \geq 0.05$ | $p \geq 0.05$ | $p \geq 0.05$ |

Finally, it is also important to recognize that intravital microscopy studies on the glomerulus are typically conducted in Munich-Wistar r
ats with superficial glomeruli, which can be routinely accessed by intravital two-photon microscopy^33, 46^. Surface cortical glomeruli are scarce in Sprague-Dawley rats^47^, and intrinsic tissue autofluorescence generally inhibits live imaging in the kidney beyond 150 μm^48^. Interestingly, the decellularization process substantially reduced the tissue autofluorescence and provided access to multiple glomeruli throughout the study, emphasizing the acquired ability to image deeper in the acellular organ and visualize these previously hidden structures. Altogether, these results indicate that an in vivo method capable of tracking microvascular integrity represents a powerful approach for studying scaffold viability and identifying ways to promote scaffold longevity and angiogenesis in bioartificial organs.

## Conclusion

Several approaches to developing a bioartificial kidney have been demonstrated, and whole organ decellularization appears to be the most promising methodology thus far^49^. One major challenge to this strategy is maintaining vascular integrity and functionality post-transplantation. Developing solutions to this problem will identify ways to promote scaffold longevity and angiogenesis in bioartificial organs. Most models used to examine the microvasculature have primarily utilized in vitro or in vivo techniques that are incapable of providing adequate spatial and temporal resolution. Thus, to further unravel mechanisms that are vital for a better understanding of this complex issue, we have employed intravital microscopy. This advanced imaging technique aids the monitoring of live cellular and subcellular events in real-time, which can provide a better understanding of the events that adversely alter scaffolds post-transplantation.

Even though the optimal conditions to achieve a decellularized whole organ have not yet been devised, current best practices have outlined SDS as a primary agent for creating acellular scaffolds. With this approach, a previously established method was used to generate whole rat kidney scaffolds. Evaluations of the decellularization process demonstrated the effective formation of acellular scaffolds and highlight the first time use of intravital fluorescent microscopy as a tool to assess scaffold generation, as well as structural and functional integrity. Specifically, these studies showed that scaffolds orthotopically transplanted into rats initially retained a reasonable degree of microvascular structure in vivo directly after transplantation. As time progressed, the scaffold succumbed to widespread coagulation and thrombosis, which would have eventually facilitated the reductions in blood flow within acellular nephrons. Simultaneously, the loss of intrinsic barriers and compensatory mechanism provided additional means to further hamper in vivo viability. Nevertheless, valuable insight into blood filtration and extravasation mechanisms were obtained and provided sites within the decellularized nephron that need to be structurally reinforced, and a possible timeline during which devastating changes occur. Again, such insight emphasizes the value of intravital microscopy to uncover subtle changes in vascular permeability, even with a limited sample size used for this study. Future studies can hopefully be conducted to address this limitation.

To ultimately create regeneration strategies suitable for clinical applications, defining optimized decellularization/recellularization parameters is essential. Thus, an experimental platform like this can improve the current understanding of events in transplanted organs. This approach can also identify ways of maintaining viable blood supply and limiting scaffold degradation in post-implantation environments. Such an understanding will support the next stage in the evolution of bioartificial organs.

## Materials and Methods

### Experimental Animals

Experiments were performed on male Sprague-Dawley and Wistar-Munich rats that ranged in weight 200-400 g (Envigo, Indianapolis, IN). All animals were given free access to standard rat chow and water throughout the study. These studies were conducted in accordance with the Institutional Animal Care and Use Committee at the School of Medicine, Wake Forest University (Winston-Salem, NC, USA), Animal Research Oversight Committee at Khalifa University of Science and Technology (Abu Dhabi, UAE), and ARRIVE guidelines. The animals were separated into the following groups (n = 4 for each group): rats in group 1 were used to supply native and decellularized kidneys respectively for DNA and SDS quantification assays; rats in group 2 were used to intravital microscopic studies of normal kidneys; and rats in groups 3, 4, 5 and 6 were used to intravital microscopic studies of transplanted acellular kidneys at the 0-hour, 12-hour, 24-hour, and 1-week marks respectively.

### Left Radical Nephrectomy

The left kidney was harvested individually by first anesthetizing an animal with inhaled isoflurane (Webster Veterinary Supply, Devens, MA, USA), 5% in oxygen, and then administering an intraperitoneal injection of 50 mg/kg of pentobarbital. Each rat was then placed on a heating pad to maintain core body temperature of roughly 30 °C. Once the animal was fully sedated, its abdomen was shaved and sanitized with Betadine Surgical Scrub (Purdue Products, Stamford, CT, USA). A midline incision was then made to expose and isolate the left kidney, and the associated renal artery, renal vein, and ureter were ligated using 4-0 silk (Fine Science Tools, Foster City, CA, USA) to extract the kidney, with intact renal arteries, veins, and ureters, from each rat. The incision was closed, and all animals were allowed to recover for roughly two weeks.

### Rat Kidney Decellularization and Sterilization

The arteries of the extracted kidneys were cannulated using PE-50 polyethylene catheter tubing (Clay Adams, Division of Becton Dickson, Parsippany, NJ) and a 14-gage cannula and then secured with a 4-0 silk suture. Kidneys were rapidly perfused with 0.5-1 ml of heparinized PBS. After heparinization, the harvested organs were suspended in PBS, and the cannulated renal artery was attached to a peristaltic pump (Cole-Palmer, Vernon Hills, IL, USA). The kidneys were then perfused via the renal artery at a 4 ml/min rate with 0.5% sodium dodecyl sulfate (Sigma-Aldrich, St. Louis, MO, USA) for a minimal perfusion time of 6 hours, followed by phosphate-buffered saline (PBS) for 24 hours. The scaffolds were then sterilized with 10.0 Ky gamma irradiation.

### DNA Quantification

DNA contents in native/control kidneys (n = 4) and decellularized kidneys (n = 4) were measured using a Qiagen DNeasy Kit (Qiagen, Valencia, CA, USA). The tissues were initially minced and stored overnight at −80°C. The next day, the tissues were lyophilized to estimate the ratios of ng DNA per mg dry tissue in each group using Quant-iT PicoGreen dsDNA assay kit (Invitrogen, Carlsbad, CA, USA) and a SpectraMax M Series Multi-Mode Microplate Reader (Molecular Devices, Sunnyvale, CA, USA) to quantify the DNA contents within the extracts.

### SDS Quantification

Decellularized scaffolds (n = 4) were first minced and then homogenized using a FastPrep 24 Tissue Homogenizer (MP Biochemicals, Santa Ana, CA, USA). The suspension was then digested with proteinase K (Omega Bio-tek, Atlanta, GA, USA) for roughly 1 h at 56 °C, after which 1 ml methylene blue solution (methylene blue 0.25 g/l, anhydrous sodium sulfate 50 g/l, concentrated sulfuric acid 10 ml/l). The samples were then extracted using chloroform and absorbance measurements were conducted at wavelength of 650 nm using the microplate reader to quantify the SDS contents within the extracts.

### Florescence Nuclear and Vascular Markers

Hoechst 33342 (Invitrogen, Carlsbad, CA, USA) and 150-kDa fluorescein isothiocyanate (FITC)-dextrans (TdB Consultancy, Uppsala, Sweden) and were used to visualize nuclei and vascular compartments, respectively, in live rats. The venous injectates were prepared by diluting 50 μl of a 20 mg/ml stock FITC-dextran solution or 30-50 μl of Hoechst in 0.5 ml of saline.

### Orthotropic Transplantation of Decellularized Rat Kidneys

After sedation, the torso of each rat was shaved and sanitized with Betadine Surgical Scrub (Purdue Products L.P., Stamford, CT). The tail vein of a sedated rat was either moistened with a warmed sheet of gauze or placed into a warm bath. A 25-gauge butterfly needle was inserted into the dilated vein tail vein and attached to a syringe containing injectates. A bolus of 0.5 ml heparinized saline was infused into the animal, and flank incisions were then created to expose the previously ligated renal artery and vein in each recipient. Note that each rat was transplanted with a biocompatible acellular scaffold that was created from its nephrectomized kidney, whereby the renal artery and vein of the decellularized kidney were anastomosed end-to-end to the remains of the recipient’s respective clamped left renal artery and vein using Micro Serrefine vascular clamps and microsurgical needles attached to10-0 silk suture thread (Fine Science Tools, Foster City, CA, USA). The ureter was also connected to a proximal portion of the renal vein using the 10-0 silk suture to maintain blood flow patterns across the organ. Other boluses of 0.25-0.5 ml of Hoechst 33342 and 150-kDa FITC-dextran molecules were introduced systemically, and then vascular clamps were removed to establish blood flow into the acellular organ. The transplanted organs were exteriorized for imaging at various measurement time points.

### Intravital Two-Photon Microscopic Assessment of Nuclear Labeling, Capillary Blood Flow, Vascular Permeability

Exteriorized native or acellular kidneys were individually positioned inside a 50 mm glass bottom dish (Willco Wells B.V., Amsterdam, The Netherlands) containing saline, set above a X60 water-immersion objective for imaging, and a heating pad was placed over the animal to regulate body temperature^28, 50^. The study was conducted using an Olympus FV1000MP multiphoton/intravital microscope (Center Valley, PA, USA) equipped with a Spectra-Physics (Santa Clara, CA, USA) MaiTai Deep See laser. The laser was tuned to excite Hoechst and FITC dyes across 770-860 nm excitation wavelengths. Images were collected with a X60 water-immersion objective and external detectors that acquired blue, green, and red emissions. The infusates were introduced via the tail vein as previously described, and static micrographs and time-series videos were collected to analyze nuclear remnants in scaffolds, changes in tissue fluorescence, and microvascular integrity/function as outlined in the literature^25, 26, 33, 41, 51^, at 0-, 12-, 24- and 1-week time points (n = 4 at each time point). ImageJ software (Fiji-ImageJ × 64, US National Institutes of Health, Bethesda, MD, USA) was used to examine changes in nuclear DNA content, microvasculature permeability and microarchitectural dimensions after transplantation, by selecting 4 regions in various renal compartments at random to measure changes in the average fluorescence intensities at the defined measurement points. The rouleaux density was determined from the number of stacked red blood cells in vascular compartments within adjacent fields divided by the vascular area, and glomerular diameters were also examined using the ImageJ software.

### Statistical Analysis

SPSS (IBM Corp, Armonk, NY, USA) was used to perform non-parametric analyses. The Kruskal-Wallis one-way analysis of variance (ANOVA) with the post hoc Dunn’s test was used to examine remnant DNA and SDS concentrations within acellular kidneys using biochemical or microscopic studies and differences in fluorescence intensities in different renal compartments between native and decellularized kidneys. This non-parametric test was also used to examine whether the degrees of dextran extravasation from the microvasculature and alterations to capillary blood flow directly after transplantation, and at 12-hour, 24-hour, and 1-week time points were significant. All variables are expressed as mean ± standard deviation, and a p-value of less than 0.05 was considered statistically significant for all evaluations.

## Acknowledgments

The authors acknowledge funding from an Institutional Research and Academic Career Development Award (IRACDA), Grant Number: NIH/NIGMS K12-GM102773, and funds from Khalifa University of Science and Technology, Grant Numbers: FSU-2020-25 and RC2-2018-022 (HEIC).

## Funding

The authors would like to thank Dr. Joao Paulo Zambon, Dr. Amanda Dillard, and Mr. Ken Grant for their support in developing the decellularization process, transplant model, and imaging protocol, respectively. The authors also wish to thank Ms. Imaan Khan for creating the graphic present in Figure 6. Finally, the authors thank Mrs. Maja Corridon, Ms. Xinyu Wang and, Dr. Adeeba Shakeel for reviewing the manuscript.

**Figure 6.**
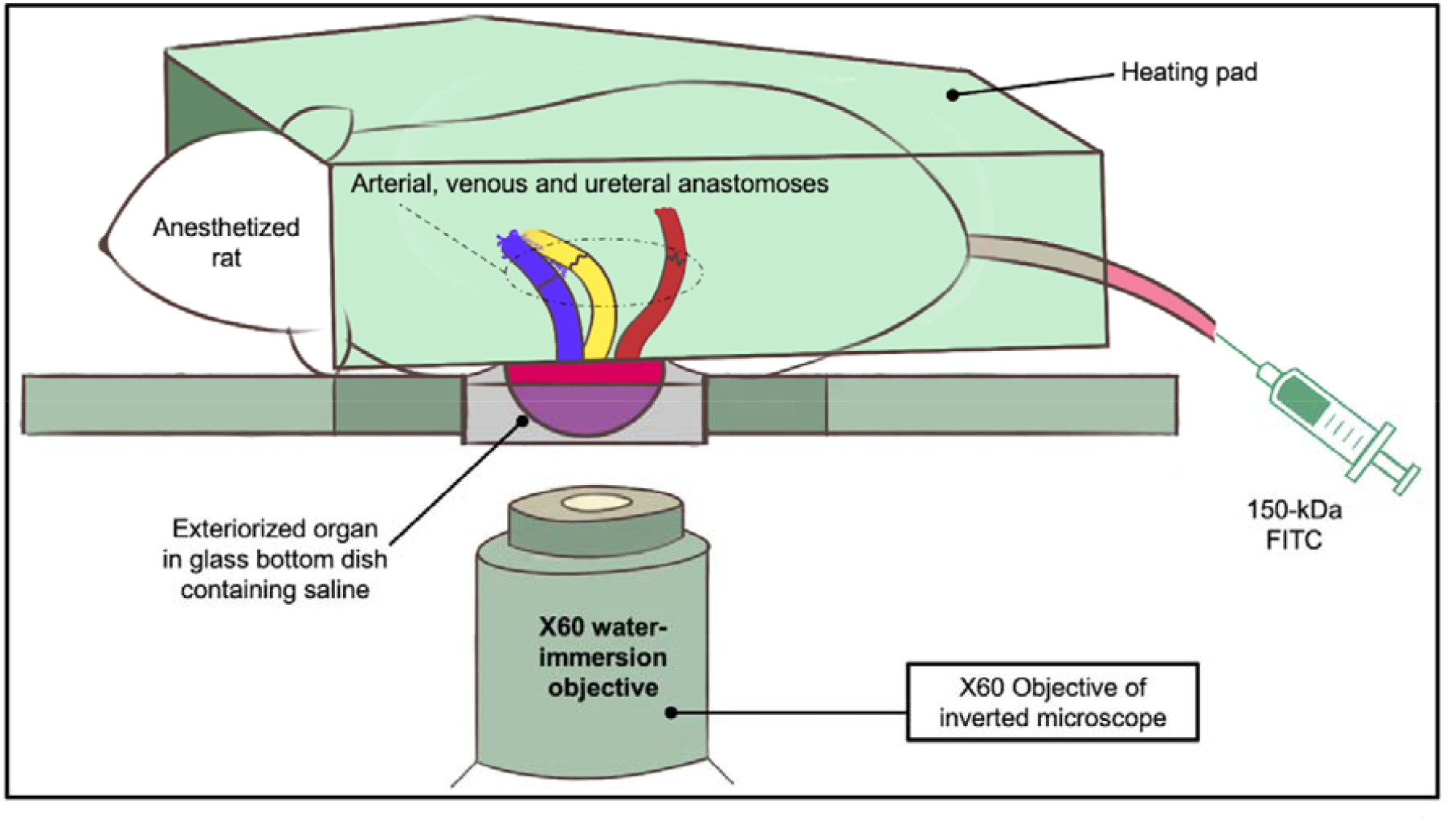
A schematic illustrating the intravital imaging process used to visualize live transplanted or native kidneys, as outlined by in the literature[28], whereby anesthetized rats with exteriorized (native or transplanted) kidneys were placed 50 mm glass bottom dish, filled with saline, and set above the stage of an inverted microscope with a Nikon ×60 1.2-NA water-immersion objective. The heating pad was then placed directly over the animal to maintain core temperature. A 25-gauge butterfly needle was inserted into the dilated vein tail vein and attached to a syringe containing injectates.

## Data Availability

Datasets generated during and/or analyzed in this study are available from the corresponding author on reasonable requests.

## Author Contributions

P.R.C. conceived and designed study and performed all the imaging and surgical procedures; P.R.C. and A.A.K. analyzed data; P.R.C. interpreted results of experiments; P.R.C. and A.A.K. prepared figures; and P.R.C. and A.A.K. prepared the manuscript and approved the final version.

## Supplementary Information

The Supplementary Material for this article can be found online at:

Video 1. Typical confinement of the 150-kDa FITC dextran vascular marker within the peritubular capillaries of native/normal rat kidney in vivo. Supplemental Video 1 available at URL: https://figshare.com/s/320b66e246b7f22454a1 DOI: 10.6084/m9.figshare.16758379

Video 2. Entry of the 150-kDa FITC dextran vascular marker into the decellularized rat nephron directly after transplantation. Supplemental Video 2 available at URL: https://figshare.com/s/0db19193096ee9ac259d DOI: 10.6084/m9.figshare.16758571

Video 3. Extravasation of the 150-kDa FITC dextran from the glomerular capillaries and its inhomogeneous and substantial translocation into decellularized proximal tubular epithelial, luminal, and interstitial compartments 24 hours after transplantation. Supplemental Video 3 available at URL: https://figshare.com/s/2f5197a256cca9e76f35 DOI: 10.6084/m9.figshare.16758778

Video 4. Significant levels of rouleaux formation and 150-kDa FITC dextran extravasation were visualized in a hypertrophied decellularized glomerulus 1-week post-transplantation. Supplemental Video 4 available at URL: https://figshare.com/s/7e60999b703a380ab1f7 DOI: 10.6084/m9.figshare.16758853

